# Cohesin is required for long-range enhancer action

**DOI:** 10.1101/2021.06.24.449812

**Authors:** Lauren Kane, Iain Williamson, Ilya M. Flyamer, Yatendra Kumar, Robert E. Hill, Laura A. Lettice, Wendy A. Bickmore

**Author notes:** Present address: Friedrich Miescher Institute for Biomedical Research, Maulbeerstrasse 66, 4058 Basel, Switzerland (Lauren Kane). Correspondence to: W.A.B: MRC Human Genetics Unit, IGC, Crewe Road, Edinburgh EH4 2XU, UK.

## Abstract

The mammalian genome is organised into topologically associating domains (TADs) that are formed through the process of cohesin-driven loop extrusion^1–3^ and whose extent is constrained at TAD boundaries by orientation-dependent CTCF binding^4–7^. The large regulatory landscapes of developmental genes frequently correspond to TADs, leading to the hypothesis that TADs and/or loop extrusion are important for enhancers to act on their cognate gene^8,9^. However, it has proven hard to interpret the consequences of experimental disruption of TADs or loop-extrusion on gene regulation^3,6,10^, in part because of the difficulty in distinguishing direct from indirect effects on enhancer-driven gene expression. By coupling acute protein degradation with synthetic activation by targeted transcription factor recruitment in mouse embryonic stem cells, here we show that cohesin, but not CTCF, is required for activation of a target gene by distant distal regulatory elements. Cohesin is not required for activation directly at the promoter or activation from an enhancer located closer to the gene. Our findings support the hypothesis that chromatin compaction mediated by cohesin-mediated loop extrusion allows for genes to be activated by regulatory elements that are located many hundreds of kilobases away in the linear genome but suggests that cohesin is dispensable for more genomically close enhancers.

## Shh as a model for enhancer function

Shh acts as a concentration-dependent morphogen during vertebrate embryonic development and the complex *Shh* regulatory domain is a paradigm for long-range enhancer regulation. Many of the described tissue-specific enhancers of *Shh* operate over large genomic distances with the regulatory landscape extending over approximately 1 Mb upstream of *Shh* (Fig. 1a).The limits of this regulatory landscape, defined using transposon-based regulatory sensors^9,11^, correspond with a TAD which contains *Shh* and all of its enhancers that have been defined so far. One of the TAD boundaries lies in an intragenic region 3’ of *Shh* whereas the other is near the *Lmbr1* promoter. The murine *Shh* regulatory landscape contains at least five CTCF binding sites (Fig. 1a), including two strongly interacting convergent sites which may form the *Shh* TAD boundaries and block loop extrusion^12^.

**Figure 1.**
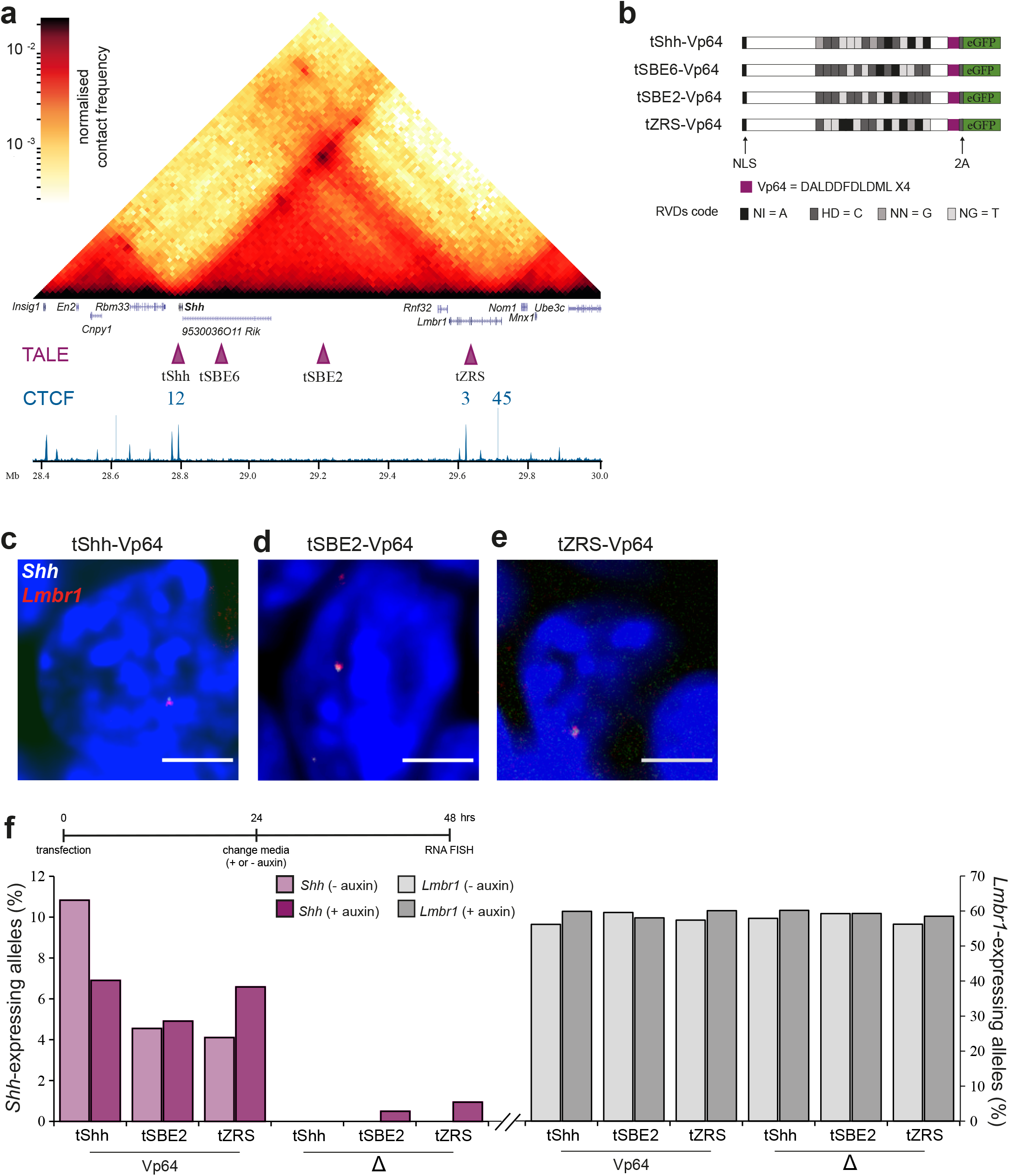
Synthetic Shh activation. (**a**) Hi-C heatmap of the *Shh* TAD from wild type mESCs at 16kb resolution. Data are from ref 28 and were created using HiGlass. Genes, positions of TALE target sequences and CTCF ChIP-seq track are shown below. Genome co-ordinates are from the mm9 assembly of the mouse genome. (**b**) Schematic of TALE-VP64 constructs used to target the *Shh* promoter (tShh-VP64), SBE62 (tSBE6-VP64) SBE2 (tSBE2-VP64) or ZRS enhancers (tZRS-VP64). NLS: nuclear localisation sequence; 2A: self-cleaving 2A peptide. Repeat variable diresidue (RVD) code is displayed at the bottom showing the one letter abbreviations for amino acids. Equivalent TALE-Δ constructs lack the Vp64 module (**c-e**) Representative images of nuclei from mESCs transfected with (**c**) tShh-VP64, (**d**) tSBE2-VP64 and (**e**) tZRS-VP64 showing RNA FISH signals for *Shh* (white) and *Lmbr1* (red). Scale bars = 5 μm. (**f**) Timecourse of TALE transfection and auxin treatment is shown above. The percent of *Shh* (left) and *Lmbr1* (right) expressing alleles in mESCs transfected with TALE-Vp64 or TALE-Δ constructs assayed by RNA FISH in the absence or presence of auxin. n = ~100 alleles. A biological replicate for these data are in Extended Data Fig. 1c.

The limb-specific ‘ZRS’, located 849 kb upstream of *Shh* in intron 5 of the widely expressed *Lmbr1,* is the most distal *Shh* enhancer. ZRS is both necessary and sufficient for *Shh* expression in the zone of polarising activity (ZPA) in distal posterior mesenchymal cells of the developing limb bud^13,14^. Increased *Shh*-ZRS colocalization, observed in the ZPA, may be consistent with a gene-enhancer interaction^15^. Large inversions that encompass the *Shh* TAD boundaries disrupt *Shh*-ZRS interactions and *Shh* regulation in limb buds^9^ but small deletions of CTCF sites at the *Shh* TAD boundaries, though disrupting TAD structure and reducing *Shh*-ZRS colocalization, do not alter the developmental pattern of *Shh* expression or cause a mutant phenotype^12^.

## Synthetic activation at long distance enhancers

Further investigation of the mechanism of long-range enhancer action and the role of cohesin-mediated loop extrusion in vivo is challenging. CTCF null mice show early embryonic lethality^16^ and conditional knockout of CTCF in the developing mouse limb results in extensive cell death^17^. Cohesin is also essential for cell proliferation^18^. However, whereas removal of cohesin in vitro does not seem to have clear immediate effects on specific gene regulation^3^, it is required for inducible gene regulation in primary haematopoietic cells^19^.

We exploited synthetic transcriptional activators to further investigate the loop extrusion model. Enhancers are activated by binding of the appropriate transcription factors (TFs) which can be mimicked by targeting of artificial TFs. Previously, we demonstrated synthetic activation of *Shh* in mouse embryonic stem cells (mESCs) using transcription activator-like (TAL) effectors (TALEs) fused to multimers of Vp16^20^. *Shh* could be activated by activator binding at the *Shh* promoter (tShh) and at the neural enhancers SBE6 (100 kb upstream) and SBE2 (410 kb upstream) (tSBE2-VP64 and tSBE2-VP64, respectively). To determine if *Shh* transcription could also be triggered by synthetic activator binding at the far end of the TAD, we designed a TALE for the ZRS (tZRS) (Fig. 1a, b).

Previously we used qRT-PCR to assay the steady state level of *Shh* mRNA induced by synthetic activators averaged across the transfected cell population^20^. To detect *Shh* nascent transcripts at a single cell/allele level, here we used RNA FISH in mESCs 48hrs after transfections with tShh-VP64, tSBE2-VP64 and tZRS-VP64 (Fig. 1c, d, e) and with control constructs lacking the activation domain (-Δ) (Extended Data Fig.1a,b). A probe set detecting *Lmbr1* nascent transcripts was used as a positive control as TALE binding was not thought to be able to affect this broadly expressed gene.

Consistent with our previous demonstration of the ability of TALE-Vp64 to activate *Shh*, *Shh* nascent RNA FISH signals were detected in mESCs transfected with tShh-VP64 (9-12% of *Shh* alleles) or tSBE2-VP64 (4-5% of alleles)(Fig. 1f, Extended Data Fig. 1b,c). Cells transfected with TALE-Δ, and non-transfected cells (ntc) showed very low signal levels. tZRS-VP64 also activated *Shh* (4% of alleles detected) indicating that *Shh* can be expressed following activator binding 850kb away. *Lmbr1* transcripts were detected at approximately 60% of alleles and these levels were similar in cells transfected with either tShh, tSBE2 or tZRS with or without fusion to Vp16 (Fig. 1f; Extended Data Fig. 1c).

## Synthetic activation in the absence of CTCF

The *Shh* TAD contains a number of CTCF binding sites (Fig. 1a) important for TAD structure but that individually are not necessary for *Shh* regulation in vivo^12^. Combinatorial deletions suggests that loss of more than one CTCF sites within the *Shh* TAD may have a more marked effect on expression^21^. Genome-wide depletion of CTCF in mESCs dramatically alters TAD insulation with rather minimal effects on ongoing gene expression^6^. However, those studies did not address where the complete loss of CTCF affects the induction of gene activation, and particularly via enhancers.

To investigate whether synthetic activation of *Shh* was dependent on CTCF we used mESCs in which the degradation of CTCF can be induced via an auxin-inducible degron (AID) (CTCF-AID)^6^. FACS (for GFP) indicated that CTCF depletion occurred as early as 6 hours after auxin addition and persisted for up to 48hrs of auxin treatment (Fig. 2a). CTCF-AID auxin-treated cells appeared to divide for at least 1-2 cell cycles, maintained a normal colony morphology and did not show significant levels of cell death during the 48 hours of auxin treatment (Extended Data Fig. 1d). Immunofluorescence indicated a very small proportion of GFP positive cells in CTCF-AID cells treated for 6 hours and so 24 hour auxin treatment, when GFP^+ve^ were completely absent, was used for subsequent experiments using CTCF-AID cells (Fig. 2b). To ensure that auxin addition did not impact TALE activity *per se*, wild type mESCs were transfected with TALE-VP64/-Δ targeting the *Shh* promoter, SBE2 and ZRS and auxin added to the media on the day after transfection for 24 hours. Targeting the *Shh* promoter or distal enhancers (SBE2/ZRS) with TALE-VP64, but not with TALE-Δ, led to activation of *Shh* expression both in the absence and presence of auxin (Fig. 1f and Extended Data Fig. 1c). *Lmbr1* expression was also consistent across all conditions and unaffected by the TALEs.

**Figure 2.**
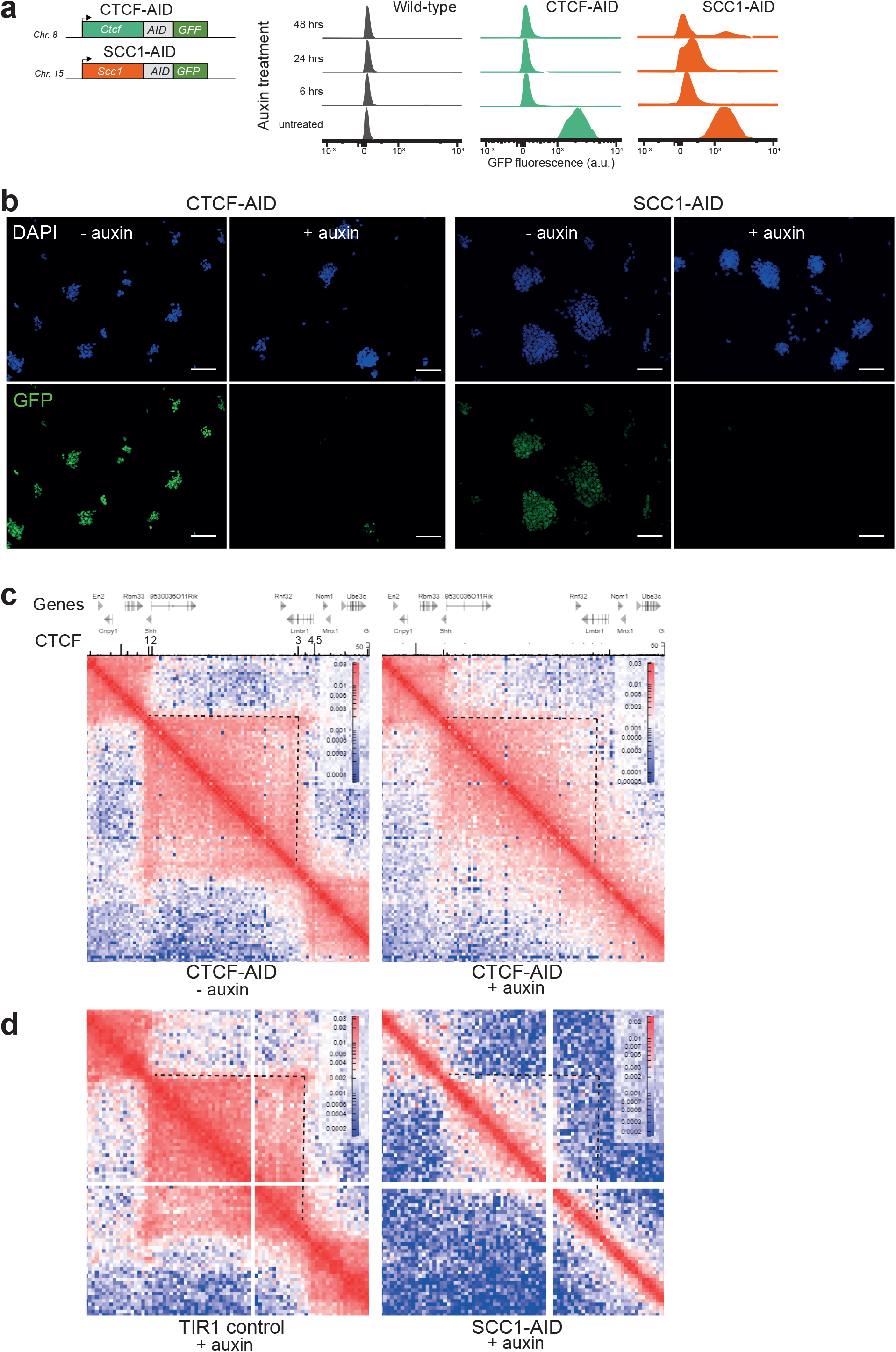
Auxin mediated degradation of CTCF and SCC1. **(a**) GFP fluorescence (a.u.) following flow cytometric analysis in wild type, CTCF-AID^6^ and SCC1-AID^22^ cells after 6, 24 and 48 hours of auxin treatment. (**b**) GFP fluorescence of untreated (-auxin) and treated (+ auxin) CTCF-AID and SCC1-AID cells after 24 and 6 hours of growth in auxin, respectively (Scale bars = 100 μm). (**c**) Hi-C heatmaps of the *Shh* TAD from untreated (-auxin) and 48 hour treated (+ auxin) CTCF-AID mESCs at 16-kb resolution (data are from ref 6). Genes (grey) and CTCF ChIP-seq (black) tracks are shown above with CTCF sites 1-5 from ref 12 indicated. CTCF ChIP-seq data shown here are from ref 6 and are from untreated (left) and auxin-treated (right) CTCF-AID cells. (**d**) Hi-C heatmaps of the *Shh* TAD from 6 hour auxin treated TIR1 control or SCC1-AID mESCs at 20-kb resolution (data are from ref 22).

Analysis of Hi-C data^6^ from auxin-treated CTCF-AID cells shows that the *Shh* TAD boundaries, and particularly that at the *Lmbr1* end, are weakened (Fig. 2c), and more inter-TAD interactions between the *Shh* and neighbouring TADs are detected in the absence of CTCF. Intra-TAD interactions were also affected by CTCF depletion, confirmed by virtual 4C display of the Hi-C data (Extended Data Fig. 2a). Using the *Shh* promoter as a viewpoint, proteolytic degradation of CTCF leads to decreased interactions of *Shh* with sequences within its own TAD and increased interactions with sequences in the adjacent *En2* containing TAD.

Given this altered 3D chromatin landscape, we tested whether CTCF depletion affected the ability to synthetically activate *Shh,* including from distal enhancers. CTCF-AID cells were transfected with tShh-VP64, tSBE2-VP64 and tZRS-VP64 and with the corresponding TALE-Δs controls. Auxin was added the day after transfection for 24 hours. *Shh* expression could still be induced in CTCF-depleted cells when targeting either the *Shh* promoter or the enhancers with TALE-VP64 (Fig. 3a, Extended Data Fig. 3a). These data suggest that activation of *Shh* expression from its distal enhancers does not require CTCF

**Figure 3.**
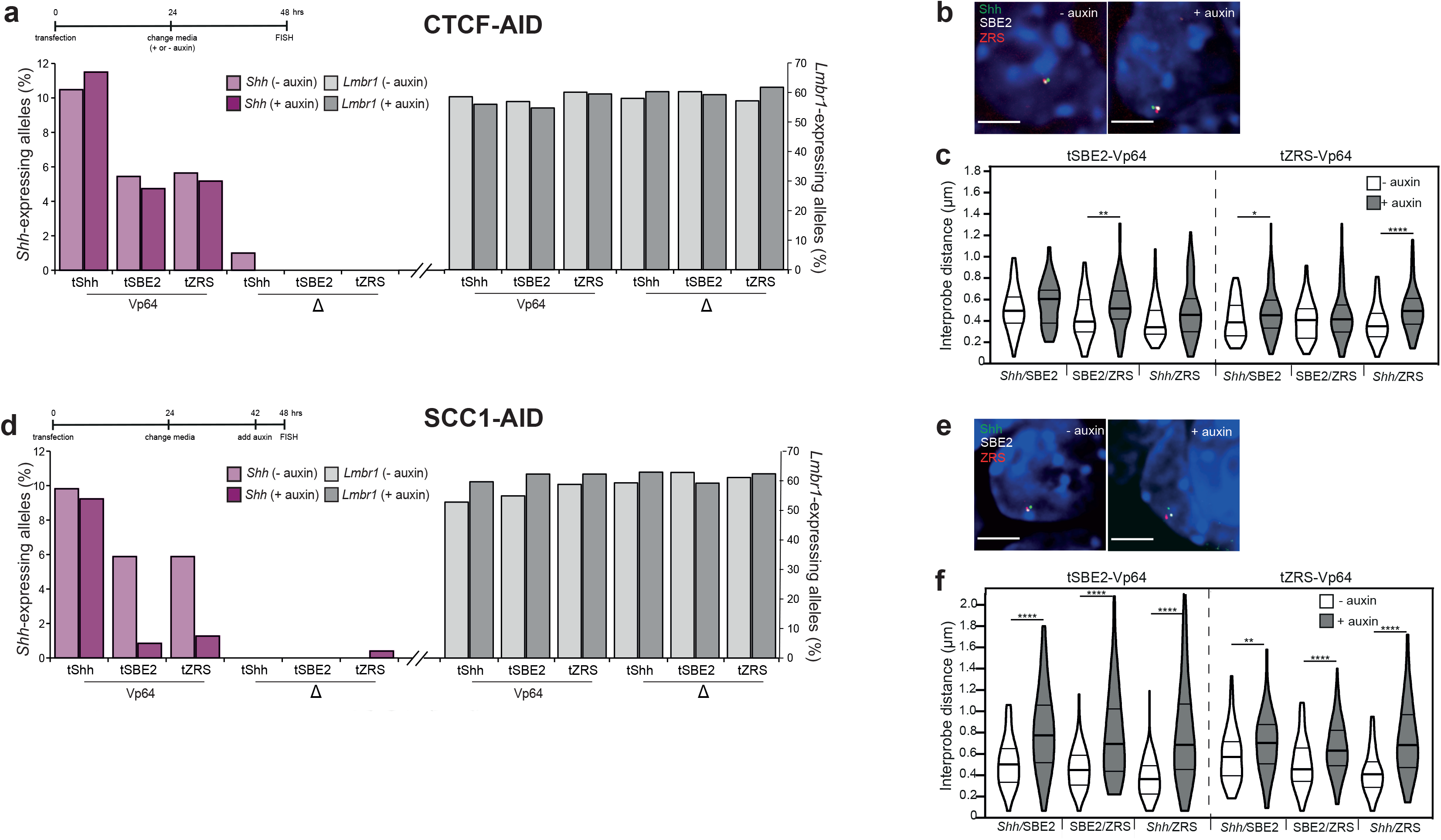
Depletion of cohesin, but not of CTCF, inhibits distal enhancer driven gene activation. (**a**) Timecourse of TALE transfection and auxin treatment is shown above. Percentage of (left axis) *Shh* and (right axis) *Lmbr1* expressing alleles, assayed by RNA FISH, in TALE-transfected CTCF-AID cells either untreated (-auxin) or treated with 24 hours of auxin (+ auxin). Cells were transfected with tShh-VP64, tSBE2-VP64 and tZRS-VP64 and equivalent TALE-Δ controls. Data shown are from one biological replicate. Data from an independent biological replicate are shown in Extended Data Fig. 3a. (**b**) Images from representative nuclei from − & + auxin CTCF-AID cells showing DNA FISH signals for *Shh*/SBE2/ZRS probes. Scale bars: 5 μm. (**c**) Violin plots showing the distribution of interprobe distances (μm) between *Shh*/SBE2, SBE2/ZRS, *Shh*/ZRS probes in tSBE2-Vp64- and tZRS-Vp64-transfected CTCF-AID cells − & + auxin. **d**) As for (**a**) but for SCC1-AID with 6 hours of auxin (+ auxin). Biological replicate for those data are in Extended Data Fig 3b. (**e**) and (**f**) As for (**b**) and (**c**) but for SCC1-AID cells. *p<0.05, **p<0.01, ****p<0.0001 (Mann-Whitney U-tests).

Consistent both with the Hi-C/ virtual 4C data (Fig. 2c, Extended Data Fig. 2a) from auxin-treated CTCF-AID cells, and with our previous analysis deleting specific CTCF sites at the *Shh* locus^12^, DNA FISH on CTCF-AID cells transfected with tSBE2-VP64 or tZRS-VP64 revealed some decompaction within the *Shh* TAD caused by CTCF loss (Fig. 3b,c and Extended Data Fig. 3c). Notably, after CTCF loss in tSBE2-VP64 transfected cells we saw no alleles where *Shh* and SBE2 were spatially co-localised (within 200nm) (Fig. 3c) despite no effect of CTCF loss on *Shh* nascent transcription by tSBE2-VP64 (Fig. 3a). This is consistent with previous observations that enhancer-gene co-localisation is not required for enhancer-driven gene-activation^20^.

## Synthetic activation from a distance is cohesin dependent

To examine effects of cohesin loss on synthetic gene activation we used auxin to acutely deplete SCC1 (RAD21) from mESCs^22^. Cohesin is required for sister chromatid cohesion during mitosis^18^ and in its absence SCC1-AID cells fail to divide, and die. FACS and immunofluorescence indicated that SCC1 depletion occurred as early as 6 hours after auxin addition (Fig. 2a, b) and we detected substantial cell death following 24 hours of auxin treatment of SCC1-AID cells (Extended Data Fig. 1d). Therefore, 6hrs of auxin treatment were used for subsequent experiments.

Genome-wide depletion of cohesin by auxin treatment of SCC1-AID is reported to erase TADs with minimal effects on steady-state gene expression^3^. Hi-C reveals a pronounced effect of SCC1 depletion on *Shh* TAD structure^22^ (Fig. 2d). Both *Shh* TAD boundaries were abrogated, and intra-TAD interactions severely depleted. Virtual 4C analysis revealed the profound loss of long-range interactions of *Shh* both within its own TAD but also with the adjacent En2 TAD (Extended Data Fig. 2b).

To assess if distal enhancers can still activate *Shh* expression in the absence of cohesin, we transfected SCC1-AID cells with TALE-VP64/-Δ proteins targeting the *Shh* promoter, SBE2 and ZRS. Similar to results for CTCF-AID, *Shh* was activated in auxin-treated SCC1-AID cells by tShh-VP64 targeting the *Shh* promoter (Fig. 3d, Extended Data Fig. 3b). *Shh* was also activated from distal sites using tSBE2-VP64 and tZRS-VP64 in SCC1-AID cells in the absence of auxin. However, synthetic *Shh* activation from these two distal sites was drastically curtailed in auxin-treated SCC1-AID cells (Fig. 3d, Extended Data Fig. 3b). *Lmbr1* expression was unaffected by the depletion of SCC1. These data suggest that SCC1/cohesin is necessary for distal activation of *Shh* by its enhancers.

As expected, given the Hi-C and virtual 4C data, in the absence of cohesin-mediated loop extrusion (SCC1 degradation), DNA FISH confirmed significant decompaction across the *Shh* TAD that was more dramatic than that seen after CTCF depletion. Significantly increased physical distances were measured between *Shh* and the distal SBE2 and ZRS enhancers (Fig. 3e,f and Extended Data Fig 3c).

## Cohesin is not required for activation from a close enhancer

These data might indicate a requirement for cohesin for activation from all enhancers, or may reflect a requirement for activation from large genomic distances. We previously demonstrated synthetic activation of *Shh* in mESCs by a TALE-Vp64 targeting SBE6 (tSBE6-Vp64)^20^, a *Shh* enhancer active in the developing brain and neural tube in vivo and neuronal progenitor cells ex vivo, and located only 100kb 5’ of *Shh* (Fig. 1a)^23^.

Surprisingly, cohesin degradation in SCC1-AID ESCs following auxin treatment did not impact on the ability of tSBE6-Vp64 to activate expression from *Shh* (Fig. 4a and Extended Data Fig. 4a). Therefore, cohesin is not essential for enhancer-driven gene activation per se, but it may be required for the function of enhancers located at relatively large genomic distances (>100kb) from their target promoter.

**Figure 4.**
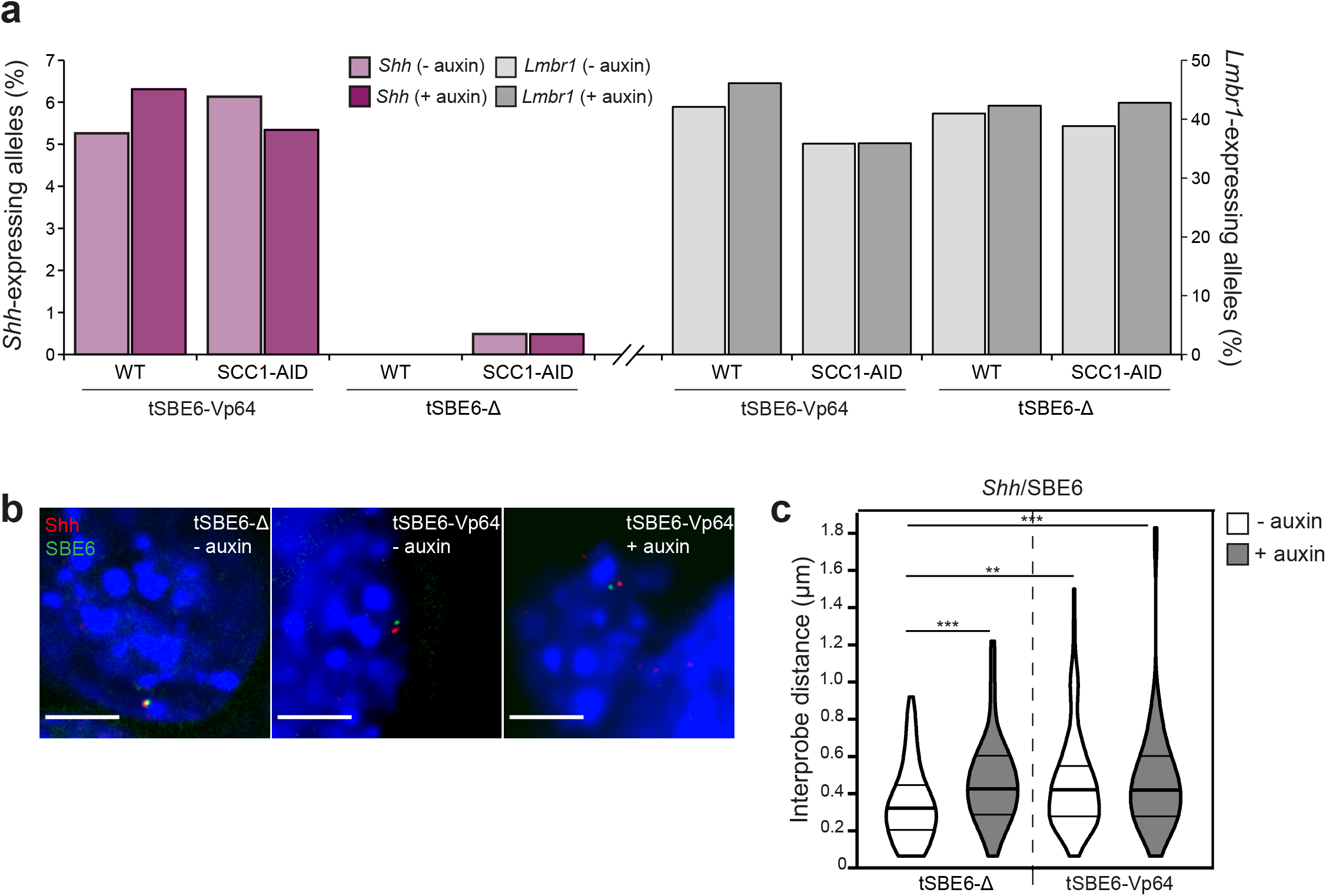
Gene activation from a close enhancer is not affected by cohesin depletion. (**a**) Percentage of (left axis) *Shh* and (right axis) *Lmbr1* expressing alleles, assayed by RNA FISH, in TALE-transfected SCC1-AID cells either untreated (-auxin) or treated with 6 hours of auxin (+ auxin). Cells were transfected with tSBE6-VP64 or tSBE6-VP64 −Δ. Data shown are from one biological replicate. Data from an independent biological replicate are shown in Extended Data Fig. 4a. (**b**) Images from representative nuclei from tSBE6-Vp64- and tSBE6-Δ-transfected SCC1-AID cells − & + auxin showing DNA FISH signals for *Shh*/SBE6 probes. Scale bars: 5 μm. (**c**) Violin plots showing the distribution of interprobe distances (μm) between *Shh*/SBE6 probes in tSBE6-Vp64- and tSBE6-Δ-transfected SCC1-AID cells − & + auxin. **p<0.01, ***p<0.001, ****p<0.0001 (Mann-Whitney U-tests).

As we have previously reported^20^, targeting of an activator to SBE6 (in the absence of auxin) led to increased spatial separation between *Shh* and this enhancer (Fig. 4b,c). Cohesin depletion also led to decompaction between *Shh* and SBE6 in the absence of activator (tSBE6-Δ). Notably, in the presence of an activator (tSBE6-VP64) cohesin depletion had no further effect on *Shh*-SBE6 distances (Fig. 4b,c and Extended Data Fig. 4b).

Our data support the observations that TAD boundaries formed by CTCF sites are not absolutely essential for an enhancer to activate its target gene located within the same TAD^12^. We find no role for either CTCF or cohesin in gene activation driven directly from the endogenous *Shh* promoter, but our data do indicate that cohesin-mediated loop extrusion is essential for activation from enhancers located at large genomic distances (>400kb) from their target gene. However, cohesin is dispensable for activation from an enhancer that is located closer (100kb) to *Shh*. This suggests that the process of loop-extrusion per se is not required for enhancer function. Rather, DNA-FISH suggests that it is the chromatin compaction brought about by loop-extrusion^24^ that may be the important factor to consider. Whereas median inter-probe distances measured at ~400kb intervals across the *Shh* TAD were modestly affected by CTCF degradation (increases of between 10 to 140nm; Figure 3c, Extended Data Fig. 3c), cohesin (SCC1) loss led to more extensive decompaction (median distance increases in the range of 130 – 330nm; Figure 3f, Extended Data Fig. 4b). In contrast, cohesin loss has no effect on chromatin decompaction between Shh and SBE6 (100kb away) when SBE6 is targeted by an activator (median distances 420nm with and without cohesin).

In contrast to a recent study examining the effect on reporter expression of genomic distance between promoters and enhancers inserted ectopically in mouse ESCs^25^, here we find no decrease in the efficiency of nascent transcription (RNA-FISH) from an endogenous promoter driven by targeted activator (Vp64) binding to different endogenous enhancer sites 100, 450 or 850kb away. One significant difference in the present study is that the activation signal has to overcome the repressive local and higher-order polycomb-mediated chromatin environment of the *Shh* locus in ESCs^26^.

The molecular mechanisms by which activating signals seeded at an enhancer transmit triggers for transcriptional activation at a distant promoter remain unclear. They might involve direct translocation of regulatory information along the chromatin fibre driven by the forces of cohesin-mediated loop extrusion, but our finding that activation from an enhancer located 100kb away from a promoter is cohesin independent argues against this model. Rather we hypothesise that loop extrusion acts to maintain the entire regulatory domain in a compact conformation^24^. This then enables either random close encounters between enhancers and promoters to initiate molecular interactions between them, or facilitates both loci to engage, for example, with the same transcriptional hub^27^. The size of this sphere of influence remains to be determined but our data examining the loss of enhancer-proximity caused by cohesin loss and the ability of targeted enhancers to activate transcription suggest that this may be <500nm, compatible with the observed distances seen between active enhancers and genes in vivo^15^.

## Acknowledgements

We thank Elphege Nora (University of San Francisco, USA) and Rob Klose (University of Oxford, UK) for their generous gifts of CTCF and SCC1-AID cell lines. We are grateful to the IGC FACs, Imaging and Technical Services facilities for their expert support. LK was supported by PhD studentship from the UK Medical Research Council (MRC). IW, YK and WAB are supported by MRC University Unit programme MC_UU_00007/2. REH and LAL are supported by MRC University Unit programme MC_UU_00007/8.

## Author Contributions

W.A.B and R.E.H conceived the study. W.A.B, LK, I.W, YK, L.A.L designed experiments. L.K and I.W performed experiments. I.M.F. analysed Hi-C data, L.K, I.W and W.A.B wrote the manuscript.

## Additional Information

Correspondence and requests for materials should be addressed to WAB.

## Methods

### Cell Lines

The mouse embryonic stem cells (mESCs) used include wild type E14 (parental line of the CTCF-AID cells provided by Elphege Nora), EN52.9.1 CTCF-AID^6^ and SCC1-AID^22^.

### Cell Culture and Transfections

Feeder-free mESCs were cultured on 0.1% gelatin-coated (Sigma G1890) Corning flasks or 10 cm dishes in GMEM BHK-21 (Gibco 21710-025) supplemented with 15% fetal calf serum (FSC; Sigma F-7524), 1000 units/mL Leukemia inhibitory factor (LIF; produced in-house), 2 mM L-glutamine, 1 mM sodium pyruvate (Sigma 58636), 5X non-essential amino acids (Sigma M7145) and 50 mM 2-β-mercaptoethanol (Gibco 31350-010). Cells were passaged at 60-90% confluence by washing with PBS, treating with trypsin (0.05% v/v; Gibco 25300-054) for ~2 minutes (mins) at 37°C and tapping flasks to detach cells. Trypsin was inactivated by addition of ten volumes of complete media and the mixture was pipetted repeatedly to generate a single-cell suspension. Cells were pelleted and plated onto gelatin-coated flasks at a density of approximately 4 × 10^4^ cells/cm^2^. Cells were maintained at 37°C with 5% CO_2_ and routinely tested for mycoplasma.

2-3 × 10^6^ mESCs were transfected with 14.5 μg of TALE plasmid and 26 μL Lipofectamine 3000 Reagent (Invitrogen L3000015) and seeded onto 0.1% gelatin-coated 10 cm dish containing an autoclaved SuperFrost Plus Adhesion glass slide. Fresh media was added to cells after 24 hours. After 48 hours of transfection, slides were washed, fixed in 4% paraformaldehyde (pFa) and permeabilised in 70% ethanol at 4°C for minimum of 24 hours (up to one week).

### Auxin-inducible degron induction

Indole-3-acetic acid (auxin) (MP Biomedicals 102037) was added to the medium either 6 (SCC1-AID) or 24 (wild type or CTCF-AID) hours prior to cell collection. 500 μM of auxin (1000X stock diluted in DMSO) was used for all experiments and stored at 4°C for up to a month or at −20°C for long-term storage.

### TALE Design and Assembly

TALEs targeting the *Shh* promoter and SBE2 had previously been designed and assembled^20^. TALE protein specific to the limb enhancer ZRS was designed using TAL Effector Nucleotide Targeter 2.0 software (https://tale-nt.cac.cornell.edu) and assembled by golden-gate assembly using a modified protocol^20, 29^. In brief, a DNA binding domain specific for a 15 nucleotide sequence was generated by the modular assembly of 4 pre-assembled multimeric TALE repeat modules (three 4-mer and one 3-mer) into a modified TALE backbone vector containing VP64-2A-eGFP. The backbone vector used for assembly of the ZRS TALE was modified to replace the ampicillin resistance cassette with spectinomycin resistance. TALE modules were picked from glycerol stocks of module library plates (Addgene 1000000034), incubated overnight at 37°C in L-broth containing 50 ng/μL ampicillin and DNA isolated using the QIAprep Spin Miniprep kit (Qiagen 27104) according to the manufacturer’s instructions. Miniprep DNA was quantified using the Quibit dsDNA broad range assay with the Quibit 4 fluorometer. TALE modules were assembled into backbone vector by setting up a 20 μL one-pot golden-gate reaction as follows: vector (100 ng), TALE modules (200ng each), 10X Tango buffer (ThermoFisher ER0451), 20 Units Esp3I (ThermoFisher ER0451), 10 Units T4 DNA ligase (New England Biosciences M0202M), 1mM ATP in ddH_2_O. Golden-gate reaction was performed on a thermal cycler ((37°C 10 mins, 16°C 10 mins x12) 36°C 15 mins, 80°C 5 mins, 4°C). Competent *E. coli* (Invitrogen LS18263012) were transformed with 5 μL of reaction.

Colonies were screened by PCR for fully assembled TALEs by setting up a 30 μL reaction as follows: single colony, 2X DreamTaq Green PCR Master mix (Thermo Scientific K1082) and 0.5 μM forward (5’GGCCAGTTGCTGAAGATCG3’) and reverse (5’CGCTACAAGATGATCATTAGTG3’) primers in ddH_2_O. Colony PCR was performed on a thermal cycler (95°C 3 mins, (95°C 30s, 55°C 30s, 72°C 120s) x30), Reaction products were run on a 1.2% agarose gel to identify positive colonies and these colonies were confirmed by Sanger sequencing. TALE-Δ constructs were made by removing the BamHI-Bg1II fragment containing VP64 from the fully assembled TALE-VP64 plasmid by restriction digest. All TALE-VP64 and TALE- Δ plasmids were miniprepped using the QIAprep Spin Miniprep kit (Qiagen 27104) according to the manufacturer’s instructions, eluted in 50 μL elution buffer, quantified using the Quibit dsDNA broad range assay with the Quibit 4 fluorometer and then stored at −20°C prior to transfection.

### RNA FISH

Custom Stellaris® RNA FISH Probes were designed against *Shh* and *Lmbr1* nascent mRNAs (pool of 48 unique 22-mer probes) using the Stellaris® RNA FISH Probe Designer (www.biosearchtech.com/stellarisdesigner (version 4.2)). Following permeabilization, slides were incubated in wash buffer (2X SSC, 10% deionised formamide) for 5 mins at room temperature. Slides were hybridized with the *Shh* and *Lmbr1* Stellaris FISH Probe set labelled with Quasar 670 and 570, respectively (Biosearch Technologies, Inc.), following the manufacturer’s instructions (www.biosearchtech.com/stellarisprotocols). RNA FISH probes were warmed to room temperature, diluted to 125 nM in Stellaris RNA FISH hybridisation buffer (#SMF-HB1-10) containing 10% formamide and hybridised to slides overnight in humidified chamber at 37°C. Slides were washed twice for 30 minutes in wash buffer at 37°C and rinsed in PBS. Slides were stained with 0.5 μg/mL DAPI and mounted using Vectashield. PBS and ddH_2_O used during RNA FISH were treated with DEPC and autoclaved to inactivate RNase enzymes. RNase free consumables were used throughout and glassware treated with RNaseZAP (Invitrogen AM9780).

### DNA FISH

Following RNA FISH, slides were re-probed for DNA FISH. Following the removal of coverslips, slides were briefly washed in PBS and then for 10 minutes in 2xSSC at 85°C followed by denaturation in 70% formamide/2xSSC at 85°C for 50 minutes before a series of alcohol washes (70% (ice-cold), 90% and 100%). 160-240 ng of biotin- and Green496-dUTP-labeled (Enzo Life Sciences) (2-colour) or biotin- and digoxigenin- and red-dUTP-labeled (Alexa Fluor™ 594-5-dUTP, Invitrogen) (4-colour) fosmid probes (Table S1) were used per slide, with 16-24 μg of mouse Cot1 DNA (Invitrogen) and 10 μg salmon sperm DNA. EtOH was added and the probe air dried. Hybridisation mix containing deionised formamide, 20 x SSC, 50% dextran sulphate and Tween 20 was added to the probes for ~1h at room temperature. The hybridisation mix containing the probes was added to the slides and the probes were hybridised to the target DNA overnight at 37°C. Following a series of washes in 2X SSC (45°C) and 0.1X SSC (60°C) slides were blocked in blocking buffer (4 x SSC, 5% Marvel) for 5 min. The following antibody dilutions were made: fluorescein anti-dig FAB fragments (Roche cat. no. 11207741910) 1:20, fluorescein anti-sheep 1:100 (Vector, cat. no. FI-6000)/ streptavadin Cy5 1:10 (Amersham, cat. no. PA45001, lot 17037668), biotinylated anti-avidin (Vector, cat. no. BA-0300, lot ZF-0415) 1:100, and streptavidin Cy5 1:10 for 3-colour detection; Texas Red avidin (Vector, cat. no. A2016) 1:500, biotinylated anti-avidin (Vector) 1:100 for 2-colour detection. Slides were incubated with antibody in a humidified chamber at 37°C for 30-60 min in the following order with 4X SSC/0.1% Tween 20 washes in between: fluorescein anti-dig, fluorescein anti-sheep, biotinylated anti-avidin, streptavidin Cy5 for 3-colour; Texas Red avidin, biotinylated anti-avidin, Texas Red avidin for 2-colour detection. Slides were treated with 1:1000 dilution of DAPI (stock 50ug/ml) for 5min before mounting in Vectashield.

**Table S1.** Fosmid Probes.

| Region | Whitehead (Sanger)<br>Name | Ensembl name | Coordinates |  | Size (bp) |
| --- | --- | --- | --- | --- | --- |
|  |  |  | Start | End |  |
| *Shh* | WI1-0574O18 | G135P64333A4 | 28754458 | 28795879 | 41421 |
| SBE6 | WI1-442E17 | G135P67311F4 | 28887686 | 28924744 | 37058 |
| SBE2 | WI1-1275C09 | G135P603171G8 | 29195832 | 29239355 | 43523 |
| ZRS | WI1-1047E14 | G135P600929F6 | 29611727 | 29653695 | 41968 |

Names are Ensembl (r 45) (http://jun2007.archive.ensembl.org/Mus_musculus/index.html). Mouse genome assembly number: NCBI m37.

### Image acquisition and deconvolution

Slides from RNA and DNA FISH were imaged using a Photometrics Coolsnap HQ2 CCD camera and a Zeiss AxioImager A1 fluorescence microscope with a Plan Apochromat 100x 1.4NA objective, a Nikon Intensilight Mercury based light source (Nikon UK Ltd, Kingston-on-Thames, UK) and Chroma #89014ET (3 colour) or #89000ET (4 colour) single excitation and emission filters (Chroma Technology Corp., Rockingham, VT) with the excitation and emission filters installed in Prior motorised filter wheels. A piezoelectrically driven objective mount (PIFOC model P-721, Physik Instrumente GmbH & Co, Karlsruhe) was used to control movement in the z dimension. Step size for z stacks was set to 0.2 μm. Hardware control and image capture were performed using Nikon Nis-Elements software (Nikon UK Ltd, Kingston-on-Thames, UK). Images were deconvolved using a calculated PSF with the constrained iterative algorithm in Volocity (PerkinElmer Inc, Waltham MA).RNA FISH signal quantification was carried out using the quantitation module of Volocity (PerkinElmer Inc, Waltham MA). Expressing alleles were calculated by segmenting the hybridisation signals and scoring each nuclei as containing 0, 1 or 2 RNA signals. DNA FISH measurements were carried out using the quantitation module of Volocity (PerkinElmer Inc, Waltham MA). For DNA FISH, only alleles with single probe signals were analysed, to eliminate the possibility of measuring sister chromatids.

### Hi-C data analysis and generation of virtual 4C profiles

Published data from ref. 6 and ref. 22 was re-analysed using the distiller pipeline (https://github.com/open2c/distiller-nf). Balanced matrices at 10 kbp were used to extract the interaction profiles of the bin containing the *Shh* promoter with the rest of the genome in all conditions. Then these profiles, and log_2_-ratio of treatment over control, were saved as bigWig files and visualised using HiGlass.

