## Extended Data Figures 1 to 4 for "Cohesin is required for long-range enhancer action"

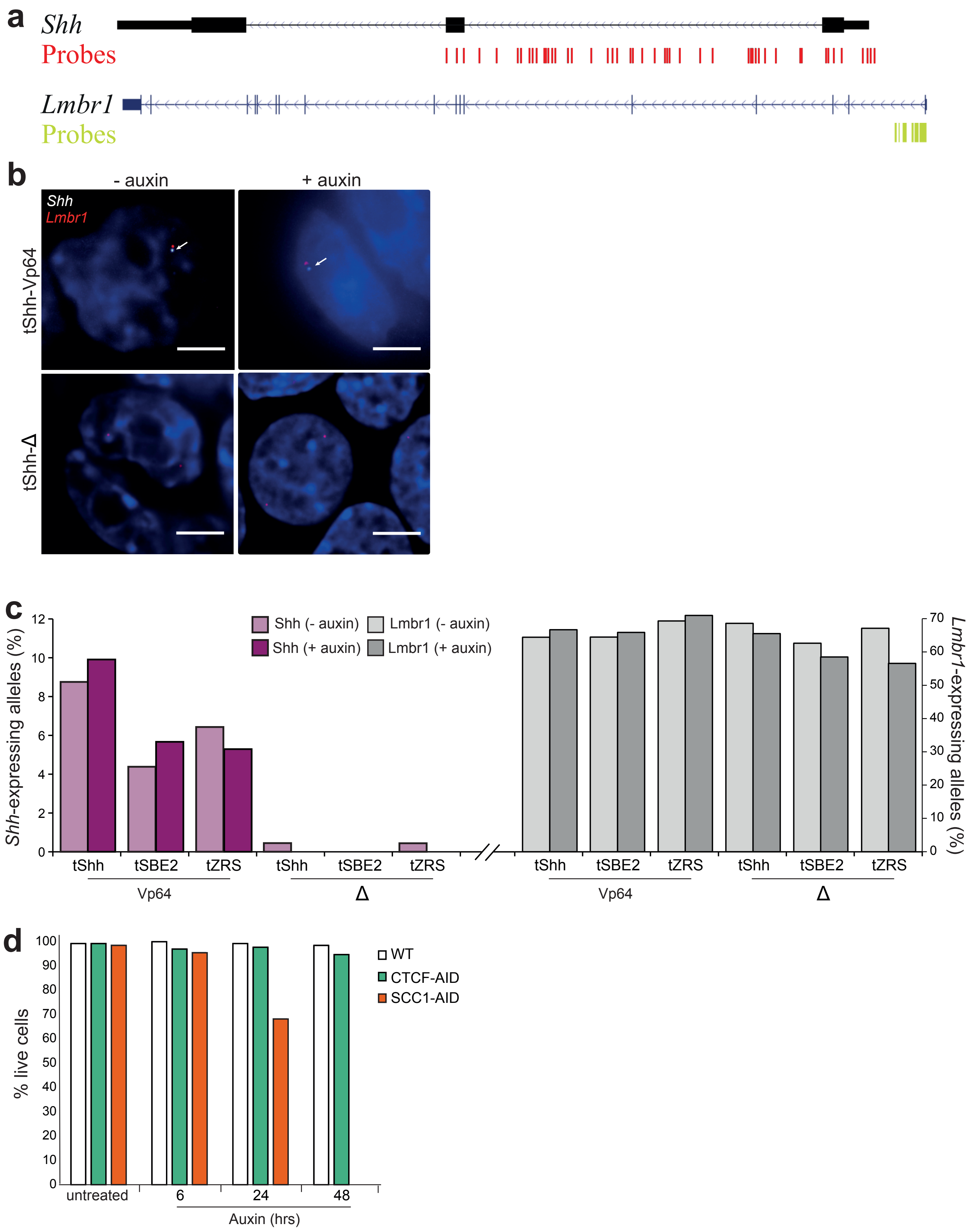


**Extended Data Figure 1. Effect of auxin treatment on mESCs**

(**a**) Schematic of *Shh* and *Lmbr1* genes showing the position of directly labelled Custom Stellaris® RNA FISH oligo probes used for RNA FISH. *Shh* probes were labelled with Quasar 670 and Lmbr1 probe were labelled with Quasar 570 (**b**) Images of representative nuclei showing RNA FISH signals for *Shh* (white) and *Lmbr1* (red) probes from wild type mESCs transfected with tShh-VP64 or tShh-Δ, and either untreated (- auxin) or treated with 24 hours of auxin (+ auxin). *Shh* RNA FISH signal is indicated by white arrow. Scale bars = 5 µm. (**c**) Quantification of the percent of (left) *Shh* (pink and red bars) and (right) *Lmbr1 – intron 1* (white and grey bars) expressing alleles in mESCs transfected with tShh-VP64, tSBE2-VP64 and tZRS-VP64 and equivalent TALE-Δ controls. Cells were either untreated (- auxin) or treated with 24 hours of auxin (+ auxin). (**d**) Quantification of live cells by DAPI staining during flow cytometry in untreated and auxin-treated wild type (WT), CTCF-AID and SCC1-AID cells after 6, 24 and 48 hours of growth in auxin.

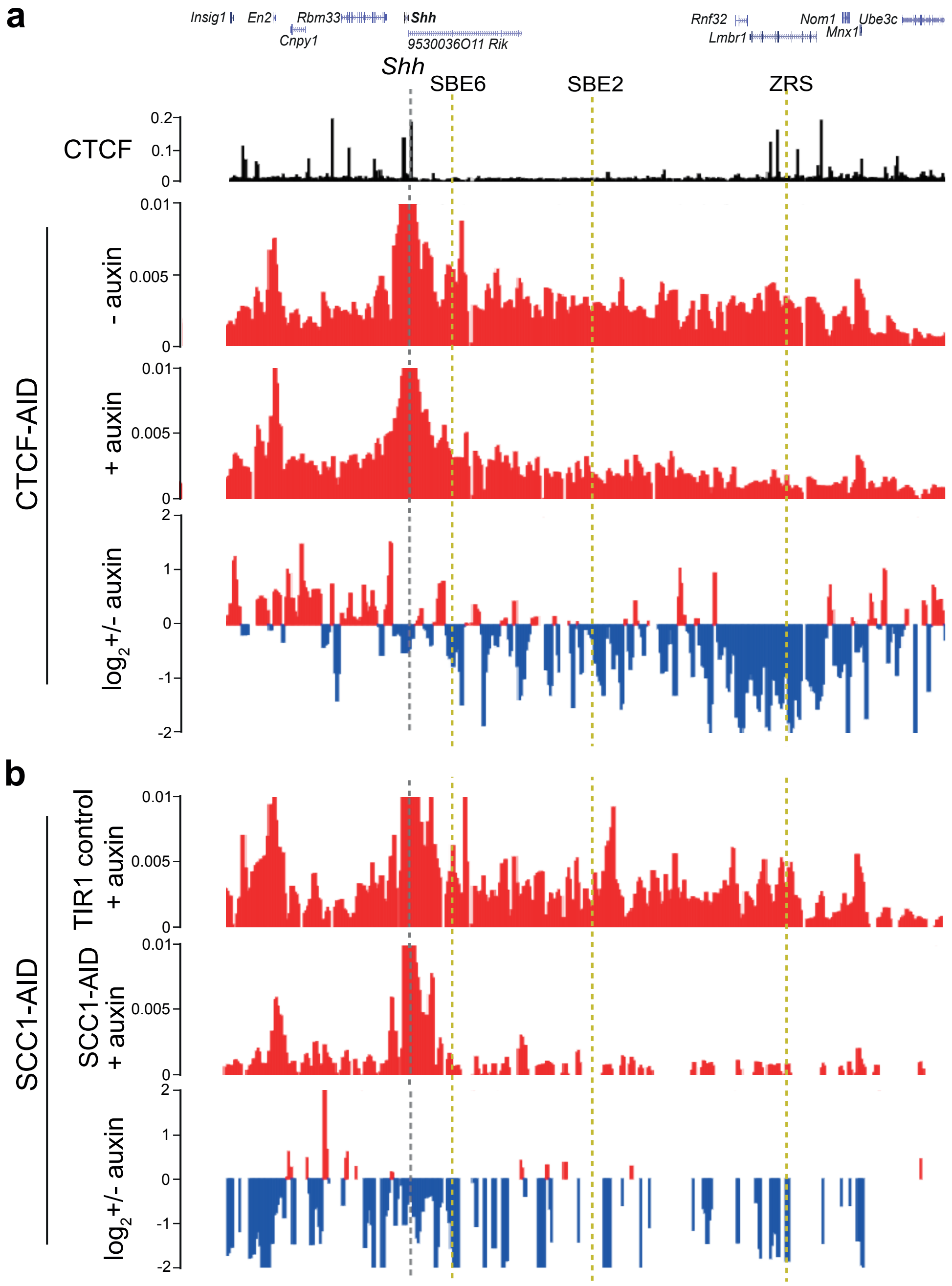


**Extended Data Figure 2. Virtual 4C following auxin mediated degradation of CTCF and SCC1**

Virtual 4C plots obtained by extracting Hi-C interactions using the *Shh* promoter as a viewpoint (grey dashed line) from untreated (- auxin) and treated (+ auxin) (**a**) CTCF-AID mESCs (data are from ref 6) or (**b**) SCC1-AID mESCs (data are from ref 22). Gene track is shown above and yellow dashed lines indicate the position of enhancers SBE6, SBE2 and ZRS. The lowest panel shows $log_2$ ratio of of data from treated vs untreated cells with gain of interactions indicated in red and loss of interactions indicated in blue.

**a**

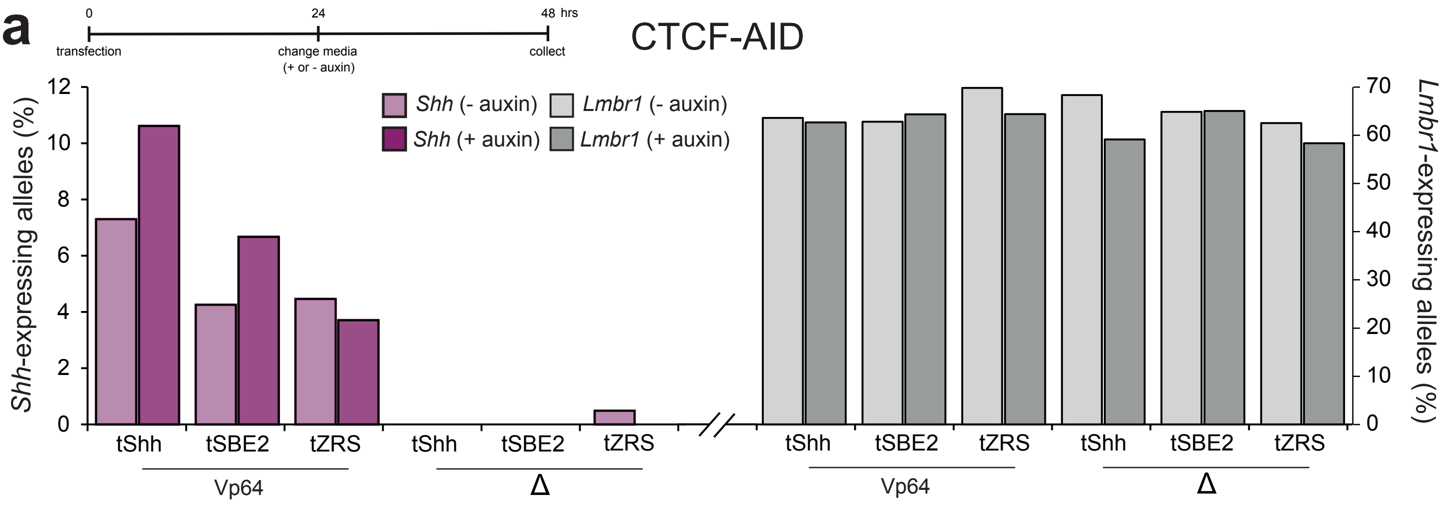


**b**

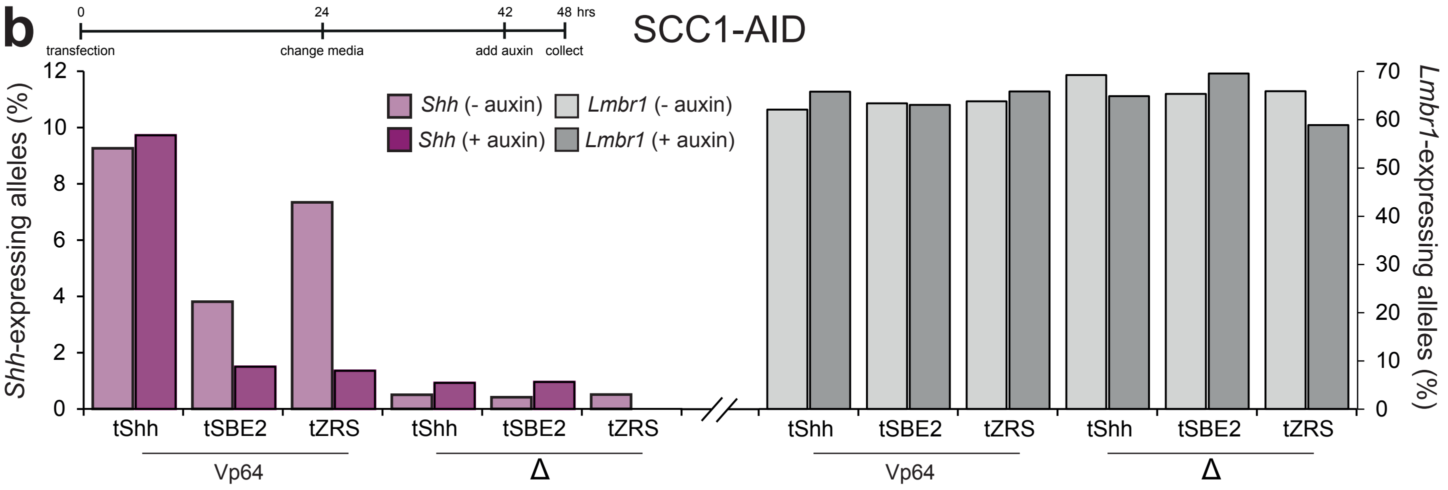


**c**

| TALE-Vp64 constructs | Probe pairs | Cell lines - or + auxin | *p* value | Median distances (μm) | Interquartile distances (μm) |
|---|---|---|---|---|---|
| **tSBE2-Vp64** | **Shh-SBE2** | **CTCF-AID –** | | 0.49 | 0.38-0.62 |
| | | **CTCF-AID +** | 0.12 | 0.61 | 0.38-0.68 |
| | | **SCC1-AID –** | | 0.50 | 0.33-0.65 |
| | | **SCC1-AID +** | **< 0.0001** | 0.77 | 0.52-1.06 |
| | **SBE2-ZRS** | **CTCF-AID –** | | 0.39 | 0.30-0.60 |
| | | **CTCF-AID +** | **0.004** | 0.52 | 0.42-0.68 |
| | | **SCC1-AID –** | | 0.45 | 0.31-0.59 |
| | | **SCC1-AID +** | **< 0.0001** | 0.69 | 0.44-1.02 |
| | **Shh-ZRS** | **CTCF-AID –** | | 0.34 | 0.28-0.50 |
| | | **CTCF-AID +** | 0.06 | 0.46 | 0.30-0.61 |
| | | **SCC1-AID –** | | 0.36 | 0.22-0.49 |
| | | **SCC1-AID +** | **< 0.0001** | 0.69 | 0.45-1.07 |
| **tZRS-Vp64** | **Shh-SBE2** | **CTCF-AID –** | | 0.39 | 0.26-0.55 |
| | | **CTCF-AID +** | **0.04** | 0.45 | 0.33-0.60 |
| | | **SCC1-AID –** | | 0.57 | 0.40-0.71 |
| | | **SCC1-AID +** | **0.003** | 0.70 | 0.51-0.88 |
| | **SBE2-ZRS** | **CTCF-AID –** | | 0.41 | 0.24-0.52 |
| | | **CTCF-AID +** | 0.43 | 0.42 | 0.30-0.55 |
| | | **SCC1-AID –** | | 0.46 | 0.34-0.66 |
| | | **SCC1-AID +** | **0.0003** | 0.63 | 0.49-0.82 |
| | **Shh-ZRS** | **CTCF-AID –** | | 0.35 | 0.25-0.47 |
| | | **CTCF-AID +** | **<0.0001** | 0.49 | 0.37-0.61 |
| | | **SCC1-AID –** | | 0.41 | 0.29-0.53 |
| | | **SCC1-AID +** | **<0.0001** | 0.68 | 0.47-0.97 |

**Extended Data Figure 3. Replicate data for effect of CTCF or cohesin depletion on distal enhancer driven gene activation**
(**a**) Quantification of the percentage of (left axis) *Shh* and (right axis) *Lmbr1* expressing alleles, assayed by RNA FISH, in TALE-transfected wild type mESCs (parental cell line used to generate the CTCF-AID cell line) and in CTCF-AID cells either untreated (- auxin) or treated with 24 hours of auxin (+ auxin). Cells were transfected with tShh-VP64, tSBE2-VP64 and tZRS-VP64 and equivalent TALE-Δ controls. Data shown are from an indepenent biological replicate of the experiment shown in Fig 3a. (**b**) As for (**a**) but for SCC1-AID with 6 hours of auxin (+ auxin). Data shown are from an indepenent biological replicate of the experiment shown in Fig 3b. **(c)** Table showing the Mann-Whitney U *p*-values for differences in inter-probe distances, for Shh-SBE2, SBE2-ZRS and Shh-ZRS probe pairs, between the data from TALE-Vp64 transfected CTCF-AID or SCC1-AID ESCs with or without the addition of auxin. Data are from Figures 3c and 3f. Median and inter-quartile distances are shown. *p*-values in bold are significant (<0.05).

**a**

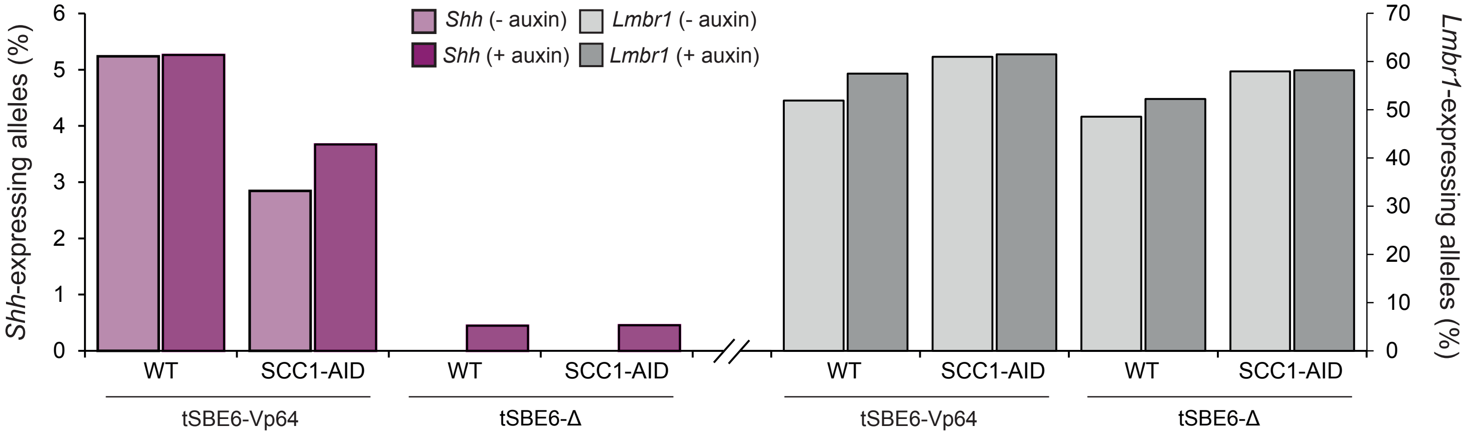


**b**

| Probe Pairs | TALE- constructs | Cell lines - or + auxin | *p* value | Median distances (μm) | Interquartile distances (μm) |
|---|---|---|---|---|---|
| **Shh-SBE6** | **tSBE6-Vp64** | **SCC1-AID –**<br>**SCC1-AID +** | 0.75 | 0.42<br>0.42 | 0.28-0.55<br>0.28-0.60 |
| | **tSBE6-Δ** | **SCC1-AID –**<br>**SCC1-AID +** | **0.0006** | 0.32<br>0.43 | 0.21-0.45<br>0.29-0.60 |

**Extended Data Figure 4. Replicate data for showing gene activation from a close enhancer is not affected by cohesin depletion**

(**a**) Percentage of (left axis) *Shh* and (right axis) *Lmbr1* expressing alleles, assayed by RNA FISH, in TALE-transfected SCC1-AID cells either untreated (- auxin) or treated with 6 hours of auxin (+ auxin). Cells were transfected with tSBE6-VP64 or tSBE6-VP64 -Δ. Data shown are from one biological replicate. Data from an independent biological replicate are shown in Fig. 4a. **(b)** Table showing the Mann-Whitney U *p*-values for differences in the Shh-SBE6 inter-probe distances between the data from SCC1-AID ESCs with or without the addition of auxin and transfected with either tSBE6-Vp64 or tSBE6-Δ. *p*-values in bold are significant (<0.05). Also shown are the median and interquartile distances of each data set. Data are from Figure 4c.
